# *TransRate*: reference free quality assessment of *de-novo* transcriptome assemblies

**DOI:** 10.1101/021626

**Authors:** Richard Smith-Unna, Chris Boursnell, Rob Patro, Julian M Hibberd, Steven Kelly

## Abstract

*TransRate* is a tool for reference-free quality assessment of *de novo* transcriptome assemblies. Using only sequenced reads as the input, *TransRate* measures the quality of individual contigs and whole assemblies, enabling assembly optimization and comparison. *TransRate* can accurately evaluate assemblies of conserved and novel RNA molecules of any kind in any species. We show that it is more accurate than comparable methods and demonstrate its use on a variety of data.

## Introduction

High-throughput sequencing of RNA has revolutionized our ability to assess the genetic basis of biological traits. For organisms that have sequenced and annotated genomes gene expression, splice-variants and mutations can be estimated from alignment of reads direct to the genome. In the absence of an appropriate reference genome, *de novo* transcriptome assembly is required.

Many *de-novo* assembly algorithms have been developed, including Trinity (Grabherr et al., 2011), Oases (Schulz et al., 2012), and SOAPdenovo-Trans (Xie et al., 2014), each of which takes a different approach to the problem. These tools provide multiple parameters and heuristics that can be modified to allow for variations in library construction, coverage depth and differences between organisms. The large parameter spaces mean that the same read data can generate substantially different assemblies both within and between assembly methods.

In addition to the considerable algorithmic flexibility, the data being assembled can be generated from multiple different RNA types. These can range from specifically amplified sub-populations of particular types of RNA, to total RNA encompassing all RNA types within the cell. Given the wide range of input data and assembly methods there is a need to be able to evaluate the quality of any *de novo* transcriptome in the absence of a known reference and identify the set of parameters, or assembly methods that best reconstruct the transcriptome from which the raw read data was generated. Moreover, there is a need to be able to identify within a given assembly the set of contigs that are well-assembled from those that are not, so that incorrect data do not influence downstream biological interpretation.

To date the majority of *de novo* transcriptome assessment methods have compared the assembled transcriptome to a known reference dataset (Lowe et al., 2014; O’Neil and Emrich, 2013). These comparative methods provide insight into the subset of known sequences that are represented within a *de novo* assembly but do not explicitly measure correctness of the assembled contigs. Furthermore, they only assess the subset of contigs that represent conserved sequences, so novel, divergent or non-coding transcripts are not assessed by these methods. Only a single reference-free transcriptome assembly evaluation tool has been published, RSEM-eval (Li et al., 2014). RSEM-eval provides an assembly likelihood given the read data, allowing the comparison of assemblies generated from the same input data. Although RSEM-eval quantifies the relative contribution that each contig makes to an overall assembly score, it is focused on assembly-level quality and does not provide descriptive statistics about the quality of each contig within an assembly.

We have developed *TransRate*, a method for evaluating the accuracy and completeness of *de novo* transcriptome assemblies that is focused on individual contigs. *TransRate* reports two key reference-free statistics: the contig score and the assembly score. The contig score measures accuracy and completeness of assembly for each individual contig by combining four independent components that are computed from the read evidence (Online Methods). These score components are designed to measure common assembly artefacts including hybrid assembly of gene families, transcript fusion (chimerism), spurious insertions, and structural abnormalities such as incompleteness, fragmentation and local mis-assembly (Figure 1). The assembly score measures accuracy and completeness of the whole assembly by combining the proportion of the read data captured with the individual contig scores. *TransRate* is free, open-source and available at http://hibberdlab.com/transrate.

**Figure 1.**
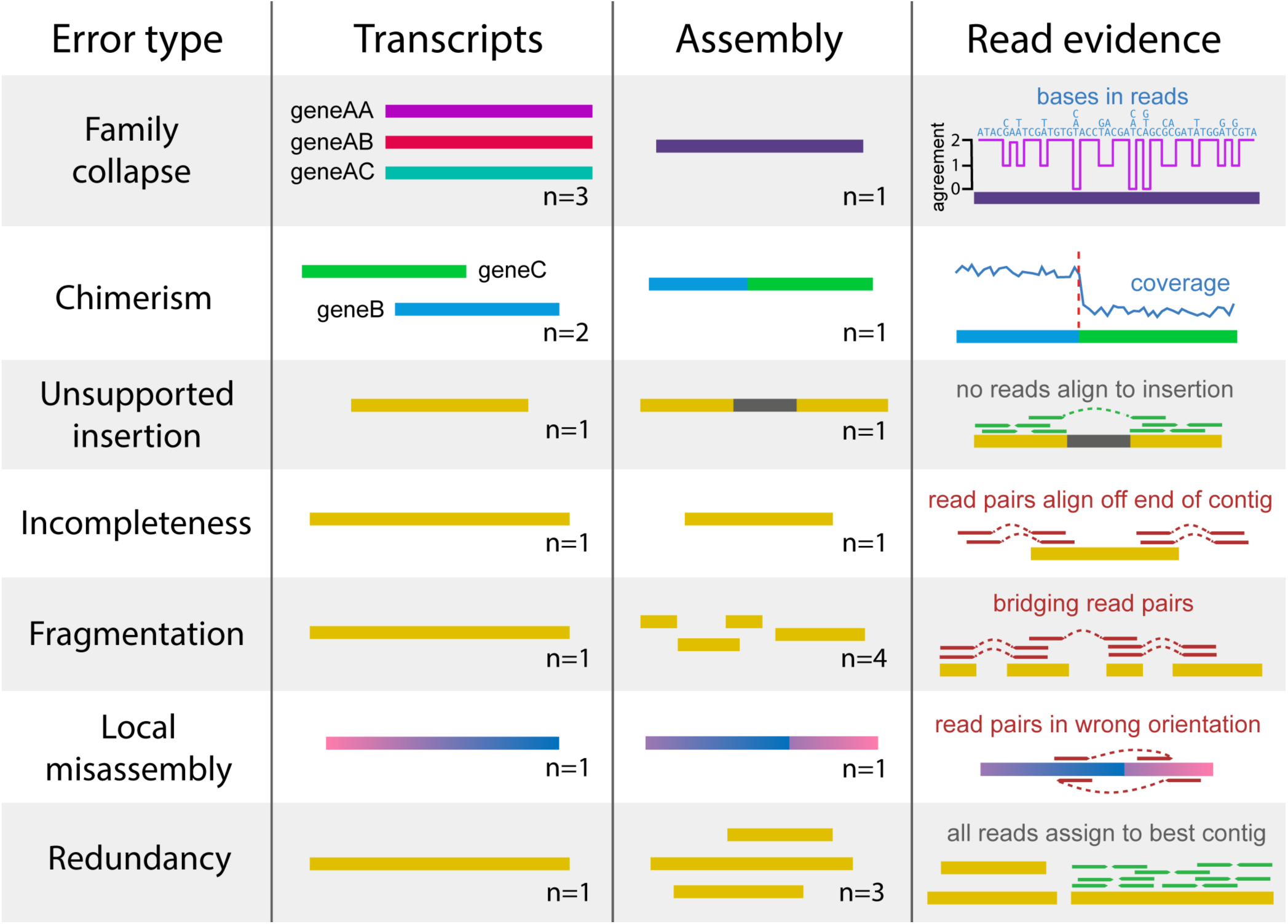
Common errors in *de-novo* transcriptome assembly, and how they can be detected using read mapping data. *Family collapse* occurs when multiple members of a gene family are assembled into a single hybrid contig. This error can be detected by measuring the extent that the nucleotides in the contig are supported by the mapped reads. *Chimerism* occurs when two or more transcripts (that may or may not be related) are concatenated together in a single contig during assembly. This can be detected when the expression levels of the transcripts differ, leading to a change-point in the read coverage along the contig. *Unsupported insertions* can be detected as bases in a contig that are unsupported by the read evidence. *Incompleteness* can be detected when reads or fragments align off the end of the contig. *Fragmentation* is detectable when read pairs bridge two contigs. *Local misassembly* encompasses various structural errors that can occur during assembly, such as inversions, usually as a result of assembler heuristics. These are detectable when both members of a read pairs align to single contig, but in manner inconsistent with the sequencing protocol. *Redundancy* occurs when a single transcript is represented by multiple overlapping contigs in an assembly. This is detectable when reads align to multiple contigs but the assignment process assigns them all to the contig that best represents the original transcript.

## Results

### Problem definition and approach

The aim of *de novo* transcriptome assembly is to accurately reconstruct the complete set of transcripts that are represented in the read data in the absence of a reference genome. There are several factors that negatively affect the accuracy of this process. These include error in the sequencing process, incomplete coverage of transcripts (due to insufficient sequencing depth), and real biological variability (such as variation in exon/intron retention, exon boundary usage, and nucleotide sequence between alleles). Errors can also originate from algorithmic simplifications such as representing the information contained in the reads as shorter words, and allowances such as allowing mis-matches that are used to reduce complexity of the assembly problem.

*TransRate* proceeds by mapping the reads to the assembled contigs, assigning multi-mapping reads probabilistically to their contig of origin, analyzing the alignments, calculating contig level metrics, integrating these contig level metrics to provide a contig score, and then combining the completeness of the assembly with the score of each contig to produce an overall assembly score (Figure 2).

**Figure 2.**
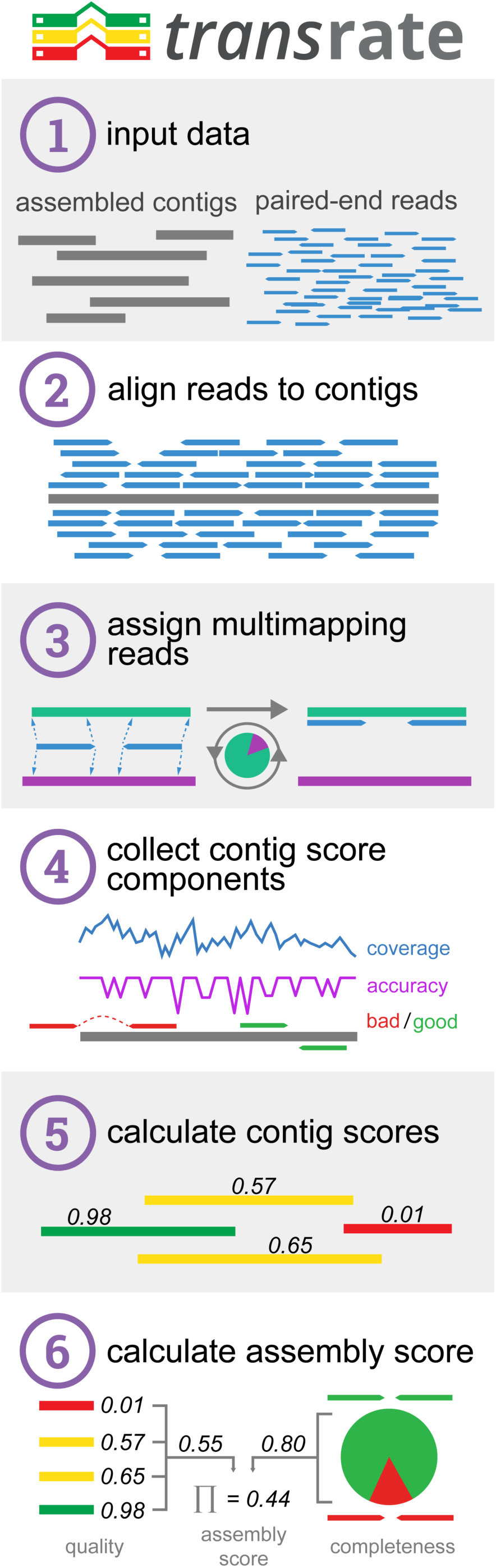
The *TransRate* workflow. (1) *TransRate* takes as input one or more *de novo* transcriptome assemblies and the paired-end reads used to generate them. (2) The reads are aligned to the contigs. (3) Multi-mapping reads are assigned to contigs based on the posterior probability that each contig was the true origin. (4) The alignments are evaluated using four independent score components. (5) The four score components are integrated to generate the *TransRate* contig score. (6) The *TransRate* assembly score is calculated from analysis of all contig scores.

### Contig assessment criteria

To calculate the *TransRate* contig score a correctly assembled contig is assumed to have the following four properties. 1) The identity of the nucleotides in the contig will accurately represent the nucleotides of the true transcript. 2) The number of nucleotides in the contig will accurately represent the number in the true transcript. 3) The order of the nucleotides in the contig will accurately represent the order in the true transcript. 4) The contig will represent a single transcript. We propose that each of these four statements can approximated through analysis of the reads that map to the assembled contigs.

*s*(*C_nuc_*) corresponds to property 1, this score measures the extent to which the nucleotides in the mapped reads are the same as those in the assembled contig. *s*(*C_cov_*) corresponds to property 2, this score measures the proportion of nucleotides in the contig that have zero coverage and thus have no supporting read data. *s*(*C_ord_*) corresponds to property 3, this score measures the extent to which the order of the bases in contig are correct by analyzing the pairing information in the mapped reads. *s*(*C_seg_*) corresponds to property 4, this score measures the probability that the coverage depth of the transcript is univariate, i.e. that it represents an assembly of a single transcript and not a hybrid/chimeric assembly of multiple transcripts expressed at different expression levels. For a detailed description of these metrics and how they are calculated see the *TransRate* contig score section.

To determine whether these four contig level metrics were independent, and thus captured different properties of each assembled contig, their performance was evaluated on a range of assemblies generated using different algorithms from multiple different species (Figure 3A). For each contig level metric the distributions of observed scores was broadly similar irrespective of species or assembly algorithm (Figure 3A). One notable exception to this observation is that the distribution of *s*(*C_cov_*) observed for rice and mouse contigs generated using SOAPdenovo-Trans was markedly different to that observed for Oases and Trinity for the same species. This reveals that the contigs generated using SOAPdenovo-Trans on this rice data contained fewer regions that had zero coverage after read mapping.

**Figure 3.**
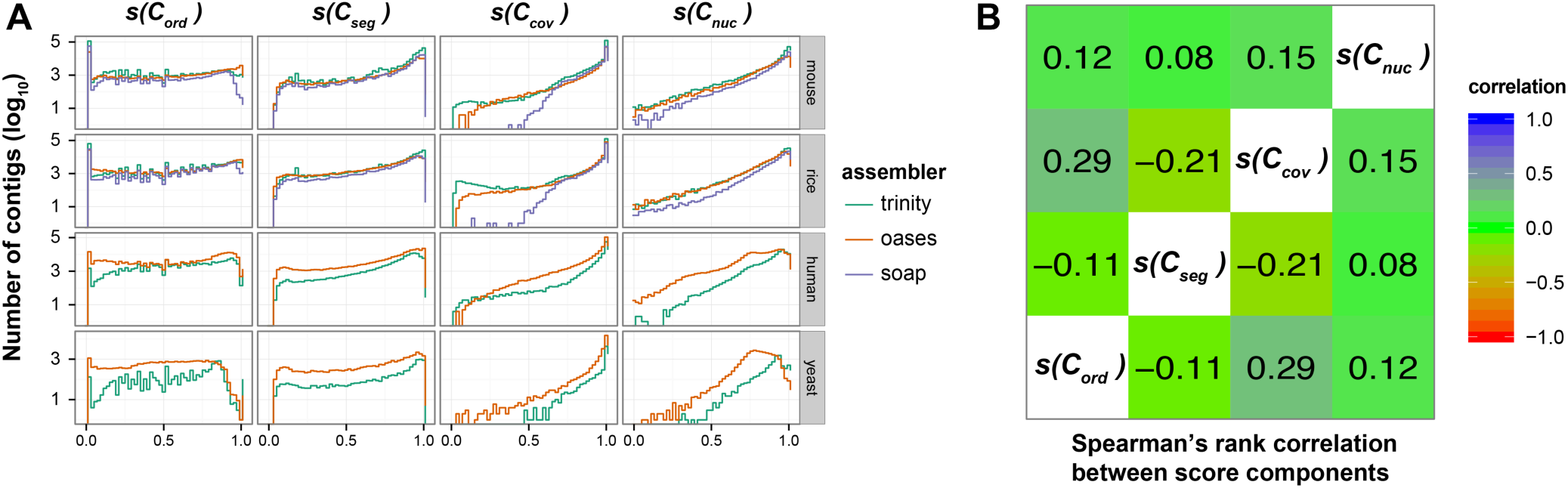
Distribution and interrelationship of contig score components. (A) Distribution of contig score components in ten different assemblies spanning four species and three different assemblers. *s*(*C_nuc_*) is the fraction of nucleotides in a contig whose sequence identity agrees with the aligned reads. *s*(*C_cov_*) is the fraction of nucleotides in a contig that have one or more mapped reads. *s*(*C_ord_*) is the fraction of reads that map to the contig in the correct orientation. *s*(*C_seg_*) is the probability that the read coverage along the length of the contig is best explained by a single Dirichlet distribution, as opposed to two or more distributions. (B) The Spearman’s rank correlation coefficient between the contig score components, averaged across all species and assemblers.

Visual inspection of the global behavior of the contig level metrics suggested that the four scores could be classified into two groups based on the density function of the observed score values. Both *s*(*C_ord_*) and *s*(*C_seg_*) produced approximately uniform distributions spanning the entire score range (Figure 3A), whereas *s*(*C_cov_*) and *s*(*C_nuc_*) produced distributions whose density increased towards higher values (Figure 3A). To determine if these visually similar distributions were correlated, and thus measured features of the assembled contigs that were inter-dependent, we analysed the pairwise Spearman’s rank correlation between the score components. This revealed that the metrics were poorly correlated (Figure 3B) and thus each provided independent assessment of the assembled contigs to which they were applied. Manual inspection of reference-based results for the 30 lowest-scoring contigs according to each score component was consistent with the individual score components accurately capturing their target properties (Supplementary file 2).

### Evaluation of the *TransRate* contig score

As the contig-level metrics provided independent evaluation of assembled contigs, we sought to determine if the product of these metrics was informative of the accuracy of assembly. To assess this, 4 million read pairs were simulated from each of the four test species (rice, mouse, human, and yeast, see Independence of score components) and assembled using SOAPdenovo-Trans with default settings. Simulated reads were used here so that the true set of transcripts was known and hence the accuracy of the assembled contigs could be assessed. The resultant assemblies were subjected to *TransRate* assessment, and the utility of the *TransRate* contig scores was assessed by comparing them to a conventional measure of contig accuracy calculated by alignment of the assembled contigs to the transcripts used to simulate the reads (see Calculation of contig accuracy). Comparison of these measures revealed that there was a strong monotonic relationship between contig accuracy and *TransRate* contig score (Figure 4A). Across all simulated datasets, the *TransRate* contig score exhibited a Spearman’s rank correlation with contig accuracy of ρ = 0.71 (Figure 4A, Supplemental file 1). For comparison we also applied RSEM-eval to the same dataset (Figure 4B). Here, the contig impact score from RSEM-eval, which measures the relative contribution of every contig to the assembly score, also showed a positive correlation with contig accuracy, however the Spearman’s rank correlation with accuracy was lower than that observed for *TransRate* (ρ = 0.36, Supplemental file 1). Non-parametric correlation measures were used here to enable unbiased comparison of *TransRate* and RSEM-eval scores, as their score distributions differ in type, location, scale and shape.

**Figure 4.**
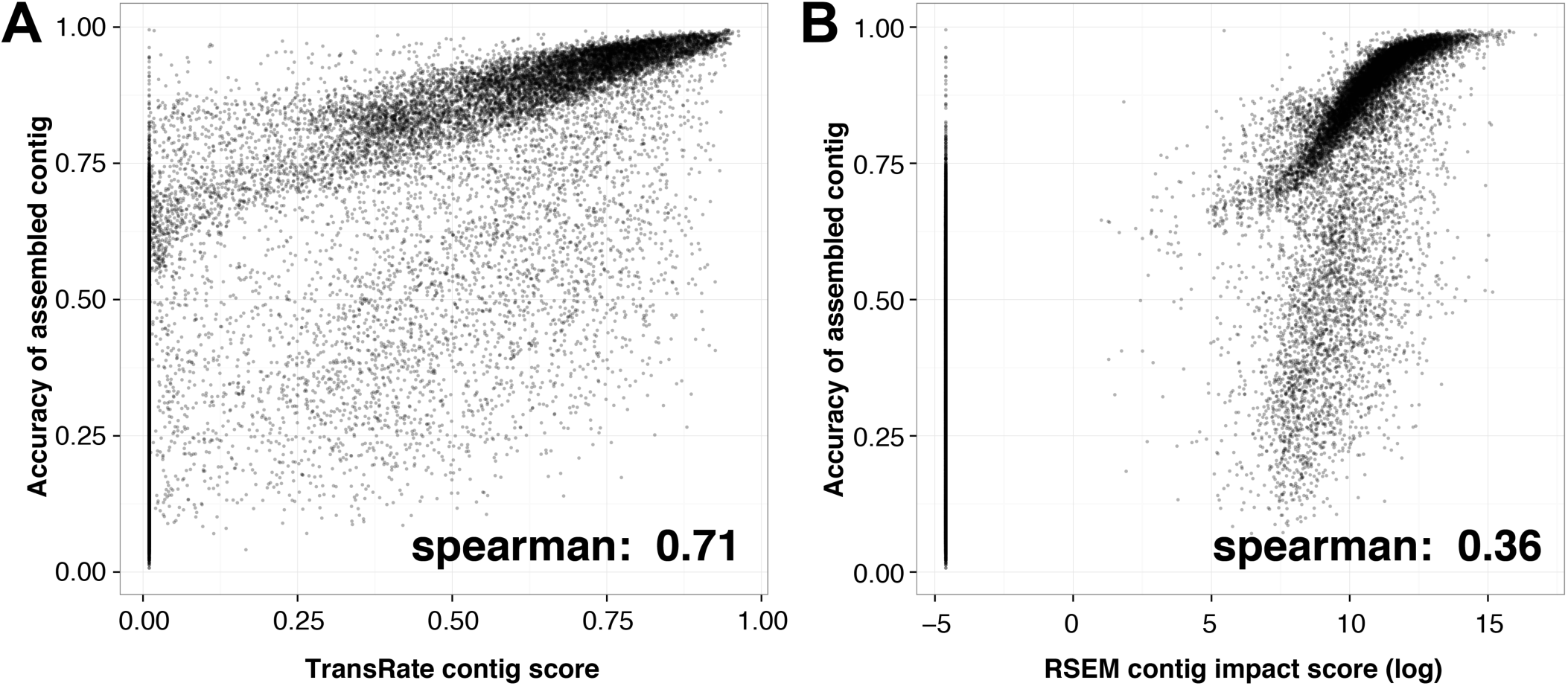
*TransRate* contig score is related to assembly accuracy. Contigs from assemblies of simulated reads from four species (rice, mouse, yeast, and human) were evaluated using *TransRate* and RSEM-eval. Reciprocal best-BLAST against the true set of transcripts was used to determine the F-score, or reference-based accuracy, of the assembled contig. Each point is a contig in an assembly, with all four assemblies on the same plot. A) Comparison between *TransRate* contig score and contig F-score. B) Comparison between RSEM-eval contig impact score and contig F-score, with contig impact scores below 0 set to the smallest positive value in the data to enable plotting.

Analysis of the interrelationship between contig scores and contig accuracy revealed that both assessment methods exhibited minimum value inflation (Figure 4A & B). Though some of these minimum value contigs comprise accurately assembled transcript sequences, they are assigned minimum score values as they fail to acquire mapped reads during the read-mapping process. This occurs due to the presence of contigs within the assembly that better represent the true contig than the contig in question and thus preferentially obtain all of the mapped reads during the probabilistic read assignment stage. This phenomenon commonly occurs when the contig in question is a substring of longer contig in the assembly. As these contigs are redundant and they would be quantified as “not expressed” in downstream expression analyses of the assemblies, both *TransRate* and RSEM-eval are justified in the assignment of minimum value scores to these contigs.

### Application of *TransRate* for relative evaluation of *de novo* assemblies from the same read data

Given that the *TransRate* contig score is strongly related to contig accuracy, we sought to develop an assembly-level score that summarised the information captured by assessment the individual contigs (Figure 4A). Here, the geometric mean of all contig scores was selected such that each contig contributed equally to the final assembly assessment. Analysis of the *TransRate* contig score distributions for assemblies generated using different assembly algorithms from different species revealed that most assemblers produced contigs that obtained a wide range of scores (Figure 5A). Some distributions also appeared to be multi-modal with overlapping populations of low and high scoring contigs (Figure 5A).

**Figure 5.**
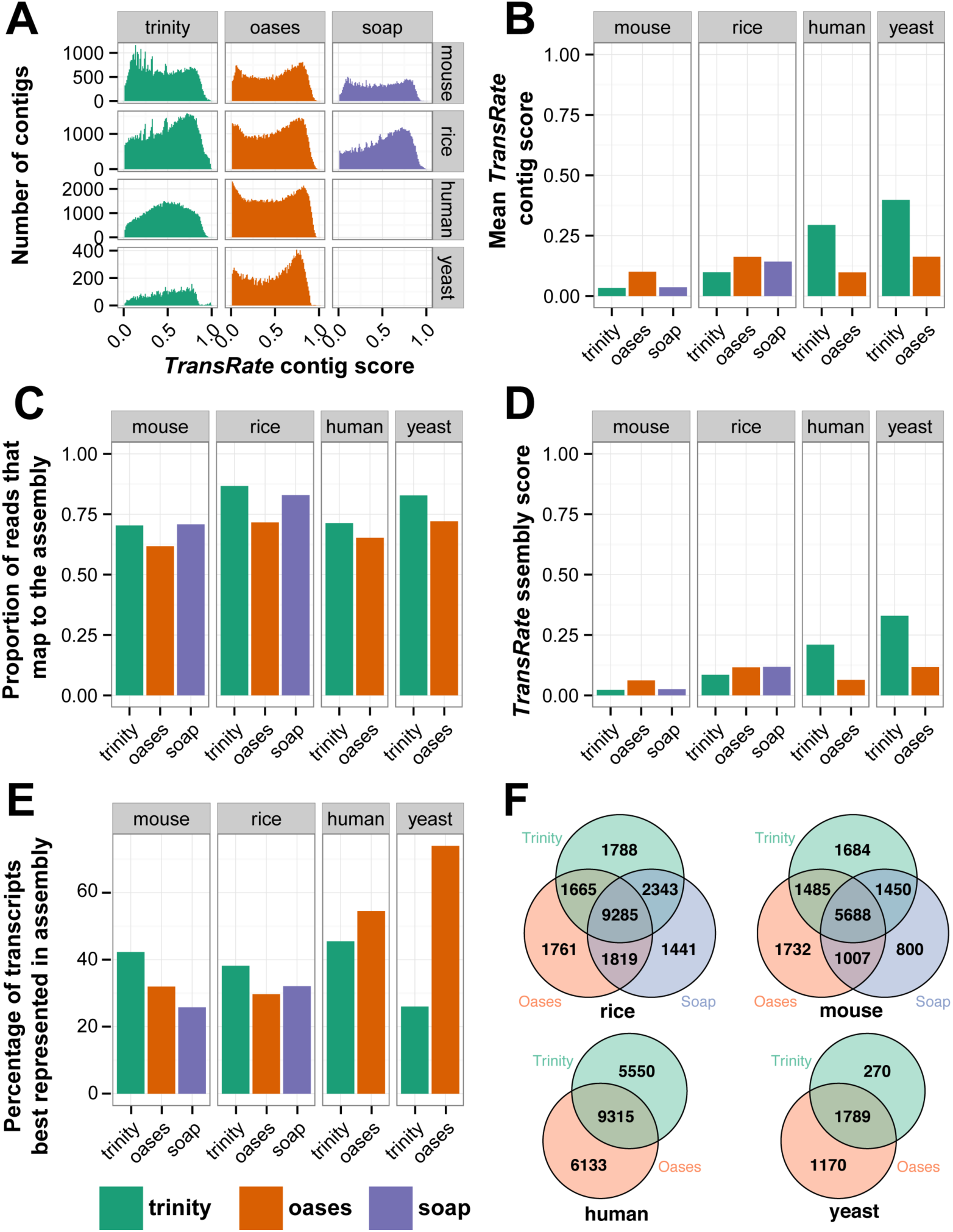
Calculation of *TransRate* assembly scores. A) Distribution of *TransRate* contig scores for the 10 representative assemblies from real data. B) Geometric mean of *TransRate* contig scores for all assemblies. C) Proportion of reads that map to each assembly. D) Final *TransRate* assembly scores for the 10 representative assemblies. E) The proportion of reference transcripts that are best assembled by individual assembly methods. F) The number of reference transcripts (identified by reciprocal best BLAST) that are assembled by each assembler.

Comparison of the geometric mean of the contig scores revealed that on different datasets, different assemblers tended to produce more accurate assemblies (Figure 5A). On average, Oases produced the highest mean contig scores for mouse and rice, while Trinity produced the highest mean contig scores for human and yeast (Figure 5B). The percentage of the input that could be mapped to these assemblies ranged from 65-85% and thus significant amounts of read data failed to be assembled by each method (Figure 5C). To provide a single assembly assessment score that combined the proportion of read data contained within the assembly and the mean accuracy of the constituent contigs we took the product of the geometric mean contig score and the proportion of reads mapping to the assembly (Figure 5D). This assembly score places equal importance on the accuracy of the assembled contigs and the proportion of the input read data that is captured by the *de novo* assembly. In an ideal scenario where all of the input reads map back to the assembled contigs with no disagreement between the reads and the assembly the assembly score will be 1. Errors in the sequencing or assembly process that cause reads to be omitted form the assembly or reads to disagree with the assembled contigs will cause the assembly score to tend towards 0.

### Further comparison of *de novo* assemblies using BLAST and *TransRate*

To demonstrate additional ways in which *TransRate* can be combined with BLAST based assessment of *de novo* transcriptome assemblies, the *de novo* assemblies was annotated using reciprocal best BLAST (bi-directional best BLAST hit) against the appropriate Ensembl reference dataset for that species. The *TransRate* scores for these contigs were compared and the proportion of transcripts that had the highest *TransRate* score for each assembly was recorded (Figure 5E). No one method consistently outperformed the others, rather the different assemblers produced the best assembly for >25% of transcripts (Figure 5E). Analysis of the total number of reference transcripts that were assembled by the different methods revealed that, though there was significant agreement between the methods, each method uniquely assembled a large number of bona fide transcripts not assembled by the other methods (Figure 5F). Taken together these analyses lend support to the idea that combining contigs from multiple assembly methods is an effective way to increase the completeness of a de novo assembled transcriptome.

### Filtration of contigs using *TransRate* contig scores

As shown in Figure 4A, 4B & 5A, many contigs within a given assembly can achieve low or minimum value scores and thus users may desire to remove them from the assembly. While *TransRate* allows the user to specify any contig score cut-off between 0 and 1 for filtration of assembled contigs, it also provides an alternative option whereby a specific contig score cut-off can be learned for any given assembly. To do this *TransRate* uses a global optimisation method to find the contig score cut-off value such that the *TransRate* assembly score function is maximised (Supplemental figure 1). This automated cut-off method is consistent with the problem definition and overall aim of *TransRate* (to assess the accuracy and completeness of a *de novo* assembled transcriptome using only the input reads) as it automatically selects the subset of contigs that maximises both accuracy and completeness.

### Comparative analysis of 155 published assemblies provides a reference for calibration and relative assessment of assembly quality

To provide a reference distribution of *TransRate* assembly scores that end-users can use to assess the relative merit of their own assemblies, *TransRate* was applied to a set of 155 published *de novo* assembled transcriptomes (Supplemental File 3). All assembled transcriptomes were downloaded from the NCBI Transcriptome Shotgun Archive (http://www.ncbi.nlm.nih.gov/genbank/tsa) and were chosen for analysis if they met the following criteria: 1) The assembly program was listed; 2) The reads were Illumina paired-end reads; 3) The published assembly contained at least 5,000 contigs. *TransRate* assembly scores for this set of published assemblies ranged tom 0.001 to 0.52 (Figure 6A, red line). Each assembly was also subject to automated assembly score optimisation producing optimised assembly scores that ranged from 0.001 to 0.6 (Figure 6A, teal line). Although some assembly scores showed little or no change following removal of low scoring transcripts, most improved when contigs below the learned cut-off were discarded (Figure 6B).

**Figure 6.**
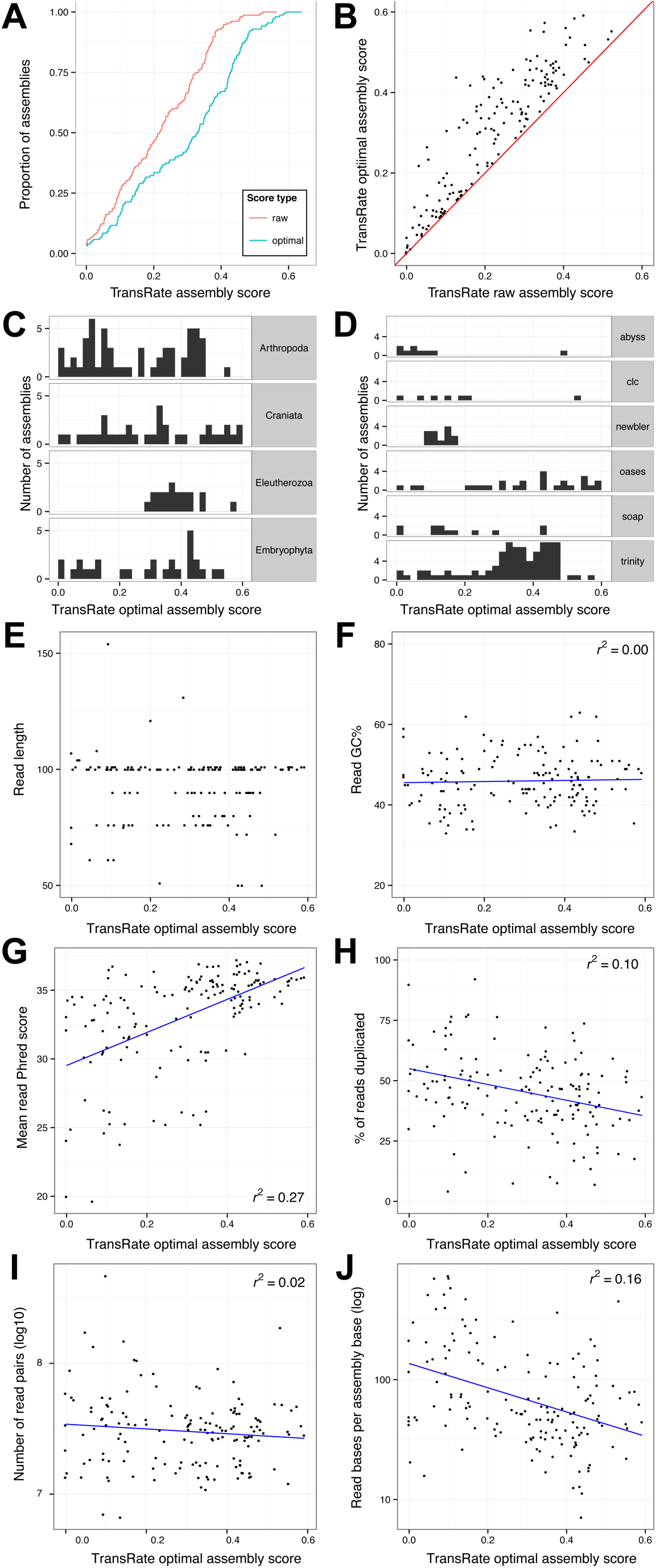
Application of *TransRate* to 155 published assemblies from the NCBI Transcriptome Shotgun Archive. 155 assemblies from the Transcriptome Shotgun Archive were analysed using transrate. The quality of the reads used to generate the assemblies were also analysed using FastQC. A) Cumulative distribution of *TransRate* raw and optimsed assembly scores for each of the 155 assemblies. B) Comparison between raw and optimised assembly score. C) Distribution of *TransRate* optimised assembly scores partitioned by taxonomic group. C) Distribution of *TransRate* optimised assembly scores partitioned by assembly method. (E-J) *TransRate* optimsied assembly scores compared to various summary statistics of the input reads: E) read length, F) read GC%, G) mean read per-base Phred score, H) percent of reads that were PCR duplicates, I) number of read pairs, and J) read bases per assembled base.

It has been suggested that the transcriptomes from certain groups of organisms may be more difficult to assemble than others (Martin and Wang, 2011). To investigate whether *TransRate* assembly scores varied for different taxa the results were analyzed according to their major phylogenetic groups (Figure 6C). For clades with more than 10 representative assemblies no association between assembly quality and taxonomic group was found (Figure 6C).

To determine if any assembler consistently produced higher *TransRate* assembly scores on end-user datasets, the performance of methods that had at least 10 assemblies was compared (Figure 6D). In this test Trinity, Oases, and SOAPdenovo-Trans all produced assemblies that spanned similar score ranges, with the highest mean score exhibited by Trinity (Figure 6D). In contrast, Newbler, Agalma and Trans-Abyss assemblies produced lower *TransRate* scores (Figure 6D). However, caution should be exercised when interpreting these results as the user-modifiable settings and post-assembly processing steps were not reported for these published assemblies. Thus the extent to which the *TransRate* assembly scores were influenced by changes in user-modifiable assembly parameters or post-assembly processing is unknown.

Given that neither assembly method nor taxonomic group produced a major effect on the *TransRate* score of an assembly we sought to determine whether the quality of the input read data was responsible for some of the variation in *TransRate* assembly scores. The read data for each assembly was analyzed using FastQC (Supplemental file 2) and the resulting read-level metrics compared to the *TransRate* assembly scores of the assemblies generated using those reads. This revealed that neither the read length nor the percentage GC of the read dataset exhibited any correlation with *TransRate* assembly score (Figure 6E & F). However, significant associations were observed for both read quality (*r*^2^ = 0.27, Figure 6G) and the level of read-duplication in the dataset (*r*^2^ = 0.1, Figure 6H). In Illumina sequencing, low read qualities are predominantly caused by errors in the sequencing process, common sources include over-clustering of the flow cell and phasing. In contrast, increases in read-duplication is caused by errors in the sample preparation stage. It occurs during the PCR amplification stage of the read library preparation and is generally caused by either conducting the library preparation from too little starting material, or by having a large variance in the fragment size such that smaller fragments become over-represented during the limited cycle PCR. While there is little correlation between the number of sequenced reads and the *TransRate* score of the assembled transcriptome (Figure 6I) there is a clear association between the relative coverage implied by those reads and the *TransRate* score (*r*^2^ = 0.16, Figure 6J). In summary, the quality of the sequence reads, the number of reads per-gene and the quality of the input cDNA library (in order of relative contribution) explain 43% of the variance in *de novo* assembly quality. Thus, the quality of the input data is more important in determining the quality of a *de novo* assembly than the choice of assembly method that is used.

## Discussion

Here we present *TransRate* a novel method for reference free assessment and filtering of *de novo* assembled transcriptomes. Our method is focused on a clear definition of an optimal *de novo* assembled transcriptome, that it should be a complete and accurate representation of the transcripts encompassed in the raw read data. *TransRate* avoids conflating assessment of *de novo* assembly quality with other criteria (such as coverage of expected reference transcript subsets) that are not equivalent to correct or complete assembly of the input reads. As the majority of published *de novo* assembled transcriptomes use Illumina paired-end sequencing, our analysis of the efficacy of *TransRate* is focused on this data type. However, the method is suitable for the analysis of other types of sequencing and thus is not restricted to use in the analysis of Illumina data.

*TransRate* is specifically designed to provide detailed insight into the quality of any *de novo* assembled transcriptome and each of its constituent contigs such that comparative analysis between assembly methods and post-assembly filtering of good and bad contigs can be performed. As *TransRate* does not use reference datasets in the evaluation of assemblies it is equally suitable for the assessment of assemblies of all types of RNA, including long non-coding RNA, mRNA, ribosomal RNA and mixed RNA samples. Moreover, given multiple assemblies generated using the same input reads, *TransRate* can also be used to determine the assembly that best represents the input read data. Thus *TransRate* could be used to help improve the performance of multiple different *de novo* transcriptome assembly algorithms.

To help end users to interpret the *TransRate* scores that they obtain for their own assemblies and place them in context of previously published assemblies, we provide a meta-analysis of 155 published *de novo* assemblies. Here, a user generated *de novo* assembly with a *TransRate* score of 0.22 (optimised score of 0.35) would be better than 50% of published *de novo* assembled transcriptomes that have been deposited in the NCBI TSA. Through detailed analysis of these 155 published assemblies we reveal that the quality of the input read data is the major factor determining the quality of any *de novo* transcriptome assembly, explaining more of the variance in quality between assemblies than quantity of read data or assembly method that is used.

## Materials and Methods

### Algorithm overview

*TransRate* is a reference-free qualitative assessment tool for the analysis of *de novo* transcriptome assemblies. *TransRate* requires one or more transcriptome assemblies and the reads used to generate those assemblies. *TransRate* aligns the reads to the assembly, processes those read alignments, and calculates contig scores using the full set of processed read alignments. *TransRate* classifies contigs into those that are well assembled and those that are poorly assembled, by learning a score cutoff from the data that maximises the overall assembly score.

### Implementation

*TransRate* is written in Ruby and C++, and makes use of the BioRuby (Goto et al., 2010) and Bamtools (Barnett et al., 2011) libraries. The source code is available at http://github.com/Blahah/transrate and is released under the open source MIT license. Binary downloads and full documentation are available at http://hibberdlab.com/transrate. The software is operated via a command line interface and can be used on OSX and Linux. *TransRate* can also be used programmatically as a Ruby gem.

### Read alignment

Reads are aligned to a given assembly using SNAP v1.0.0 (Zaharia et al., 2011). Alignments are reported up to a maximum edit distance of 30. Up to 10 multiple alignments are reported per read where available (-omax 1⊘), up to a maximum edit distance of 5 from the best-scoring alignment (-om 5). Exploration within an edit distance of 5 from each alignment is allowed for the calculation of MAPQ scores (-D 5). BAM-format alignments produced by SNAP are processed by Salmon v0.4 (Patro et al., 2014) so that multi-mapping reads are assigned to a single contig based on the posterior probability that the reads come from that contig.

### *TransRate* contig scores

*TransRate* outputs scores for every contig. Here, an assembly consists of a set of contigs *C* derived from a set of reads 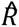. Reads are aligned and assigned to contigs such that *R_I_* is the set of reads assigned to *C_I_*. We propose that a correctly assembled contig derived from a *de novo* transcriptome assembly will have the following four intuitive properties.

1. **The identity of the nucleotides in the contig will accurately represent the nucleotides of the true transcript** *s*(*C_nuc_*). This score measures the extent to which the nucleotides in the mapped reads are the same as those in the assembled contig. If the mapped reads do not support the nucleotides of the contig then this likely because: A) The non-supportive reads should map to a different contig or a contig that is not represented in the assembly (a similar gene family variant, alternative allele, or other similarly encoded gene), or B) the assembled sequence is incorrect. In the case of the former, a missing contig (i.e. one that is not assembled) will negatively affect the score of the contig to which its reads incorrectly map. Though the contig to which they map may be correctly assembled, the negative score for this contig can be justified as the incorrectly mapped reads will render the abundance estimate of the assembled contig invalid. In the case of the latter, disagreement between the reads and the contig must be due to mis-assembly. To ensure that stochastic read errors that result in disagreement between a read and a contig do not affect the overall score for that contig support for an alternative nucleotide sequence needs to be provided by multiple reads, (see below).
2. **The number of nucleotides in the contig will accurately represent the number in the true transcript** *s*(*C_cov_*). This score measures the proportion of nucleotides in the contig that have zero coverage and thus have no supporting read data. If there are nucleotides in the contig that are not covered by any reads (regardless of the agreement between the reads and the sequence of the contig) then this should negatively impact on the contig score.
3. **The order of the nucleotides in the contig will accurately represent the order in the true transcript** *s*(*C_ord_*). This score measures the extent to which the order of the bases in contig are correct by analyzing the pairing information in the mapped reads. Here, if the orientation of the mapped reads does not conform to an expected mapping estimated from an analysis of a sub-sample of mapped read pairs then these incorrectly mapping reads will negatively affect the contig score. Similarly, if the contig could have been extended, i.e. there are read-pairs that map such that one read is present near a terminus of the contig and its pair is not mapped and would be expected to map beyond the scope of the contig, then such cases indicate that the contig does not use all of the available reads and thus is incompletely assembled.
4. **The contig will represent a single transcript** *s*(*C_seg_*). This score measures the probability that the coverage depth of the transcript is univariate, i.e. that it represents an assembly of a single transcript and not a hybrid/chimeric assembly of multiple transcripts expressed at different expression levels. Here the per-nucleotide coverage depth of the contig must be best modelled by a single Dirichlet distribution (described below). If the contig is better modelled by the product of two or more Dirichlet distributions then this indicates that two or more contigs with different transcript abundances have been erroneously assembled together.

The *TransRate* contig score is the product of the scores for each of these properties using the aligned reads as evidence. These four properties is evaluated as follows.

### Calculation of *s*(*C_nuc_*)

The alignment edit distance is used to quantify the extent to which the contig sequence is correct. The alignment edit distance is the number of changes that must be made to the sequence of a read in order for it to perfectly match the contig sequence. Here the edit distance of an aligned read r_i, j_ ∈ R_i_ is denoted as *e*_*r*_*i j*__ and the set of reads that cover nucleotide *k*(*k ∈* [1,*n*]) as ϱ*k*. The maximum possible edit distance for an alignment is limited by the read alignment algorithm (described in the Read alignment section above) and is denoted as 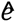 The support for the contig provided by the reads is then evaluated as 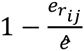 for each r_i_ ∈ ϱ*k*, and the mean of all support values is used to calculate *s*(*C_nuc_*)

### Calculation of *s*(*C_cov_*)

This score is evaluated as the fraction of nucleotides in the contig that receive at least one mapped read irrespective of the agreement between the read and the contig.

### Calculation of *s*(*C_ord_*)

The pairing information of the mapped reads is used to evaluate this score. To determine the parameters of the read library preparation a randomly selected sub-sample of 1% of all mapped read pairs are analyzed. From these alignments the orientation of the paired end reads is determined and the mean and standard deviation of the fragment size is inferred. All read pair alignments are then classified according to whether they are plausible given the estimated parameters of the library preparation and assuming that the assembled contig is correct. A read pair is considered correct if the following criteria are met: (a) both reads in the pair align to the same contig, (b) the relative orientation of the reads in the pair is consistent with the inferred library preparation parameters, (c) the relative position of the reads is consistent with the mean and standard deviation of the inferred fragment size. *s*(*C_nuc_*) is then evaluated as the proportion of all mapped read pairs that are correct.

### Calculation of *s* (*C_seg_*)

The per-nucleotide read coverage data is used to evaluate this score. To evaluate the probability that the contig originates from a single transcript (i.e. it is not chimeric) a Bayesian segmentation analysis of the per-nucleotide coverage depth is performed. For a correctly assembled contig it is assumed that the distribution of per-nucleotide coverage values in that contig is best described by a single Dirichlet distribution. i.e. all nucleotides in the same transcript should have the same expression level and thus should be best modelled as a stochastic sample from a single distribution. In contrast, a contig that is a chimera derived from concatenation of two or more transcripts will have per-nucleotide coverage values that are best described by two or more different Dirichlet distributions. The probability that the distribution of per-nucleotide read coverage values comes from a single Dirichlet distribution is evaluated using a Bayesian segmentation algorithm previously developed for analysis of changes in nucleotide composition (Liu and Lawrence, 1999). To facilitate the use of this method, the per-nucleotide coverage along the contig is encoded as a sequence of symbols in an unordered alphabet by taking log_2_ of the read depth rounded to the nearest integer. As the probability will be a value between 0 and 1, this probability is used directly as *s*(*C_seg_*).

### TransRate assembly score

The aim of the *TransRate* assembly score is to provide insight into the accuracy and completeness of any given assembly. Thus the assembly score weights equally a summary statistic of the *TransRate* contig scores and the proportion of the input reads that are contained within this assembly. We note here that alternative methods for summarizing contig scores (such as weighting contig scores by their expression level) would produce different results. However, such schemes would not be consistent with the problem definition and aim of *TransRate*: to assess the accuracy and completeness of a *de novo* assembled transcriptome using only the input reads. This score assumes that an ideal assembly will contain a set of contigs that represent unique and complete transcripts to which all of the reads used to assemble those transcripts can be mapped. The *TransRate* assembly score (*T*) is evaluated as the geometric mean of the mean contig score and the proportion of read pairs that map to the assembly such that

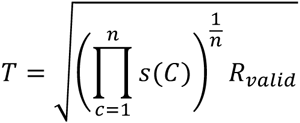

Where

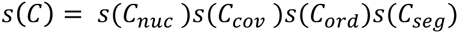

### Analysis of assemblies generated from real reads

To demonstrate the utility *TransRate* contig and assembly scores using real data, *TransRate* was applied to publicly available benchmark assemblies from two previous analyses (Davidson and Oshlack, 2014; Xie et al., 2014). One set comprised different assemblies generated for rice (*Oryza sativa*) and mouse (*Mus musculus*) using the Oases, Trinity, and SOAPdenovo-Trans assemblers (Xie et al., 2014). The other set comprised assemblies for human (*Homo sapiens*) and yeast (*Saccharomyces cerevisiae*) that had been assembled with Oases and Trinity (Davidson and Oshlack, 2014). These assemblies were chosen as they have previously been independently used in benchmark comparisons and each of the species has a completed annotated reference genome available. In all cases, *TransRate* was run with the published reads and the published assembly as input.

### Independence of score components

Correlation between the contig score components was measured for the assemblies from real data. To prevent larger assemblies from biasing the results, 5,000 contigs were sampled at random from each assembly. These contigs were used to calculate a Spearman’s rank correlation coefficient using R version 3.1.1 (R Core Team, 2014). The correlation between any two score components was taken as the mean of the correlation across all datasets.

### Identification of reconstructed reference transcripts

The full set of coding and non-coding transcripts for each species were downloaded from Ensembl Genomes version 25 (ftp://ftp.ensemblgenomes.org/pub/release-25/). Assembled contigs were then identified by BLAST searching the reference dataset for the corresponding species using bidirectional blastn local alignment with an e-value cutoff of 10^−5^ (BLAST+ version 2.2.29 (Camacho et al., 2009)). Only reciprocal best hits were retained for further analysis.

### Assembly from simulated read data

For each species, a total of 10 million mRNA molecules were simulated from the full set of annotated mRNAs from the Ensembl reference with exponentially distributed expression values using the flux simulator v1.2.1 (Griebel et al., 2012). mRNA molecules were uniform-randomly fragmented and then size-selected to a mean of 400 nucleotides and standard deviation of 50 nucleotides. From the resulting fragments, 4 million pairs of 100bp reads were simulated using the default error profile included in flux-simulator. An assembly was generated from these simulated reads using SOAPdenovo-Trans with default parameters.

### Calculation of contig accuracy

Accuracy was calculated by comparing contigs assembled from simulated data to the set of transcripts from which the read data were simulated. Reciprocal best BLAST hits were identified and the accuracy of each contig assembled from simulated read data was evaluated as the contig F-score where

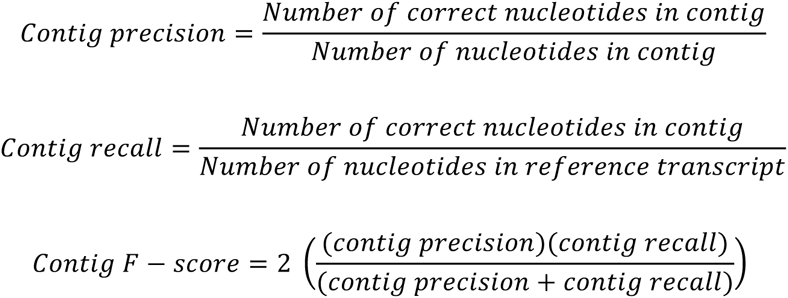

Spearman’s rank correlation coefficient between the contig F-score and the *TransRate* contig score was calculated using R version 3.1.1. The same contigs were also subject to analysis using RSEM-eval and the relationship between contig impact score and contig F-score analyzed using the same method.

### Constructing a benchmark dataset of *TransRate* scores

A survey of the range of assembly scores for published *de novo* transcriptome assemblies was conducted by analyzing a sub-set of transcriptome assemblies from the Transcriptome Shotgun Archive (http://www.ncbi.nlm.nih.gov/genbank/tsa). *De novo* assembled transcriptomes were used in this analysis only if paired-end reads were provided, the assembler and species were named in the metadata, and the assembly contained at least 5,000 contigs (*TransRate* has no minimum or maximum contig requirements but a minimum number of 5,000 was imposed to ensure sufficient raw data was available for analysis). For each of these test datasets, the assembly and reads were downloaded. *TransRate* was run on all assemblies and FastQC version 2.3 (Andrews, 2010) was used to evaluate the quality of the read datasets.

### Analysis implementation

The experimental and analysis code, implemented in Ruby and R, are available under an MIT license at https://github.com/Blahah/transrate-paper.

## Acknowledgements

The authors thank the *TransRate* user community for testing, bug reports and feedback. In particular, we thank Matt MacManes for meticulous testing. RSU was funded by The Millennium Seed Bank (Royal Botanical Gardens, Kew). SK was supported the Bill and Melinda Gates Foundation and UKAID as part of the C_4_ rice project. CB was supported by a grant to JMH from the Biotechnology and Biological Sciences Research Council, the Department for International Development and (through a grant to BBSRC) the Bill & Melinda Gates Foundation, under the Sustainable Crop Production Research for International Development programme, a joint initiative with the Department of Biotechnology of the Government of India’s Ministry of Science and Technology.

## Supplemental file legends

**Supplemental file 1.** CSV. Spearman’s rank correlation between the contig F-score, the *TransRate* contig score, and the RSEM-eval contig impact score, for all contigs in simulated assemblies.

**Supplemental file 2.** PDF. Visualisation of alignment evidence for lowest scoring 30 contigs according to each score component in Trinity assembly of yeast.

**Supplemental file 2.** CSV. *TransRate* results and read analysis summary statistics for Transcriptome Shotgun Database assemblies.

**Supplemental Figure 1.** PNG. Example of the global optimisation method used to learn the automated contig score cut-off. Here, the optimal cut-off is the value that maximises the *TransRate* assembly score.

## References

Andrews, S., 2010. FastQC: a quality control tool for high throughput sequence data [WWW Document]. URL http://www.bioinformatics.babraham.ac.uk/projects/fastqc

Barnett, D.W., Garrison, E.K., Quinlan, A.R., Strömberg, M.P., Marth, G.T., 2011. BamTools: a C++ API and toolkit for analyzing and managing BAM files. Bioinformatics 27, 1691–1692. doi:10.1093/bioinformatics/btr174

Camacho, C., Coulouris, G., Avagyan, V., Ma, N., Papadopoulos, J., Bealer, K., Madden, T.L., 2009. BLAST+: architecture and applications. BMC Bioinformatics 10, 421. doi:10.1186/1471-2105-10-421

Davidson, N.M., Oshlack, A., 2014. Corset: enabling differential gene expression analysis for de novo assembled transcriptomes. Genome Biol. 15, 410. doi:10.1186/s13059-014-0410-6

Goto, N., Prins, P., Nakao, M., Bonnal, R., Aerts, J., Katayama, T., 2010. BioRuby: bioinformatics software for the Ruby programming language. Bioinformatics 26, 2617–2619. doi:10.1093/bioinformatics/btq475

Grabherr, M.G., Haas, B.J., Yassour, M., Levin, J.Z., Thompson, D.A., Amit, I., Adiconis, X., Fan, L., Raychowdhury, R., Zeng, Q., Chen, Z., Mauceli, E., Hacohen, N., Gnirke, A., Rhind, N., Palma, F. di, Birren, B.W., Nusbaum, C., Lindblad-Toh, K., Friedman, N., Regev, A., 2011. Full-length transcriptome assembly from RNA-Seq data without a reference genome. Nat. Biotechnol. 29, 644–652. doi:10.1038/nbt.1883

Griebel, T., Zacher, B., Ribeca, P., Raineri, E., Lacroix, V., Guigó, R., Sammeth, M., 2012. Modelling and simulating generic RNA-Seq experiments with the flux simulator. Nucleic Acids Res. 40, 10073–10083. doi:10.1093/nar/gks666

Li, B., Fillmore, N., Bai, Y., Collins, M., Thomson, J.A., Stewart, R., Dewey, C.N., 2014. Evaluation of de novo transcriptome assemblies from RNA-Seq data. Genome Biol. 15, 553. doi:10.1186/s13059-014-0553-5

Liu, J.S., Lawrence, C.E., 1999. Bayesian inference on biopolymer models. Bioinformatics 15, 38–52. doi:10.1093/bioinformatics/15.1.38

Lowe, E.K., Swalla, B.J., Brown, C.T., 2014. Evaluating a lightweight transcriptome assembly pipeline on two closely related ascidian species. PeerJ Prepr. 505. doi:10.7287/peerj.preprints.505v1

Martin, J.A., Wang, Z., 2011. Next-generation transcriptome assembly. Nat. Rev. Genet. 12, 671–682. doi:10.1038/nrg3068

O’Neil, S.T., Emrich, S.J., 2013. Assessing De Novo transcriptome assembly metrics for consistency and utility. BMC Genomics 14, 465. doi:10.1186/1471-2164-14-465

Patro, R., Mount, S.M., Kingsford, C., 2014. Sailfish enables alignment-free isoform quantification from RNA-seq reads using lightweight algorithms. Nat. Biotechnol. 32, 462–464. doi:10.1038/nbt.2862

R Core Team, 2014. R: A language and environment for statistical computing. R Foundation for Statistical Computing, Vienna, Austria.

Schulz, M.H., Zerbino, D.R., Vingron, M., Birney, E., 2012. Oases: robust de novo RNA-seq assembly across the dynamic range of expression levels. Bioinformatics 28, 1086–1092. doi:10.1093/bioinformatics/bts094

Xie, Y., Wu, G., Tang, J., Luo, R., Patterson, J., Liu, S., Huang, W., He, G., Gu, S., Li, S., Zhou, X., Lam, T.-W., Li, Y., Xu, X., Wong, G.K.-S., Wang, J., 2014. SOAPdenovo-Trans: de novo transcriptome assembly with short RNA-Seq reads. Bioinformatics btu077. doi:10.1093/bioinformatics/btu077

Zaharia, M., Bolosky, W.J., Curtis, K., Fox, A., Patterson, D., Shenker, S., Stoica, I., Karp, R.M., Sittler, T., 2011. Faster and More Accurate Sequence Alignment with SNAP. ArXiv11115572 Cs Q-Bio.

